# Long-term consequences of alcohol use in early adolescent mice: focus on neuroadaptations in GR, CRF and BDNF

**DOI:** 10.1101/2021.11.26.470081

**Authors:** P. Sampedro-Piquero, R.D. Moreno-Fernández, A. Begega, M. López, L.J. Santín

## Abstract

Our aim was to assess the cognitive and emotional state, as well as related-changes in the glucocorticoid receptor (GR), the corticotropin-releasing factor (CRF) and the brain-derived neurotrophic factor (BDNF) expression of adolescent C57BL/6J male mice after a five-week two-bottle choice protocol (postnatal day (pd) 21 to pd52). Additionally, we wanted to analyse whether the behavioural and neurobiological effects observed in late adolescence (pd62) lasted until adulthood (pd84). Behavioural testing revealed that alcohol during early adolescence increased anxiety-like and compulsive-related behaviours, which was maintained in adulthood. Concerning cognition, working memory was only altered in late adolescent mice, whereas object location test performance was impaired in both ages. In contrast, novel object recognition remained unaltered. Immunohistochemical analysis showed that alcohol during adolescence diminished BDNF+ cells in the cingulate cortex, the hippocampal CA1 layer and the central amygdala. Regarding hypothalamic-pituitary-adrenal axis (HPA) functioning, alcohol abuse increased the GR and CRF expression in the hypothalamic paraventricular nucleus and the central amygdala. Besides this, GR density was also higher in the prelimbic cortex and the basolateral amygdala, regardless of the animals’ age. Our findings suggest that adolescent alcohol exposure led to long-term behavioural alterations, along with changes in BDNF, GR and CRF expression in limbic brain areas involved in stress response, emotional regulation, and cognition.

## 1. INTRODUCTION

Risk-taking behaviours, such as excessive alcohol consumption, are common during human adolescence [1, 2] in most western countries. This increased propensity to consume alcohol is also enhanced by the fact that adolescents tend to be significantly less sensitive to alcohol’s negative reinforcing properties than adults, including *hangover* symptoms and increased negative affect during alcohol withdrawal [3, 4]. Nevertheless, lasting negative emotional and cognitive states following alcohol consumption has been described in both adolescent humans and laboratory rodents in tasks such as the novelty suppressed feeding test or the forced swimming test [5, 6], among others. Further, the potential role of alcohol use as a risk factor for adult psychiatric disorders and alcohol dependence is remarkable [7, 8].

These negative consequences are due in part to the alcohol impact on brain mechanisms and signalling systems, whose maturation mainly happens during adolescence, when the vulnerability of the developing brain to the toxic effects of alcohol is higher [9, 10]. Thus, early alcohol exposure can produce impairments in the brain structure and function, resulting in several behavioural and cognitive deficits [11]. For instance, studies have revealed persistent alcohol-induced neurobiological changes within the prefrontal cortex (PFC), the hippocampus and the amygdala integrally involved in governing diverse emotional states [12, 13, 14]. Besides this, these brain regions regulate the hypothalamic-pituitary-adrenal (HPA) axis activity, which is also a critical mechanism contributing to protracted alcoholism [15, 16]. Clinical and experimental studies in both humans and rodents have shown that both acute and chronic alcohol consumption, as well as alcohol withdrawal, enhanced plasma glucocorticoids (GCs) and decreased GCs receptors’ (GR) availability [17, 18]. Interestingly, although the relationships between HPA axis activity, craving, and behavioural performance during early abstinence have been documented, little is known on such a relationship during protracted abstinence [19, 20, 21, 22]. The early abstinence period is associated with a decrease in plasmatic GCs concentration, as opposed to a brain regional GCs increase, particularly in the PFC, likely involving genomic effects of GCs [23]. Hence, it seems that the transition from positive to negative reinforcement in alcohol dependence is driven by a dysregulated HPA axis function, in which the role of GRs and GCs remains to be investigated.

On the other hand, alcohol also alters the level and function of neurotrophic factors such as the brain-derived neurotrophic factor (BDNF) [24, 25]. It is expressed throughout the nervous system, with the highest presence in the cortex and hippocampus [26]. BDNF has been linked to pathways that negatively regulate the adverse actions of ethanol as well as promote positive neuroplasticity and anxiolytic-related behaviours [27, 28, 29]. There is a large body of evidence suggesting that glucocorticoid and neurotrophin activities are closely connected [30]. Hence, a sustained hypercorticoid state diminishes BDNF expression in brain regions such as the hippocampus, attenuating neuroplasticity and its inhibitory tone on the HPA axis [31]. This suggests a vicious circle through which stress, by inhibiting BDNF, elevates GCs levels, affecting a variety of higher-order brain functions [32].

In this context, the present study has aimed to study the consequences of alcohol use during adolescence in emotional behaviours, cognitive responses and stress-related markers (BDNF, GR and CRF) in different brain regions in male C57BL/6J mice. We have focused on the medial prefrontal cortex, the paraventricular nucleus, the bed nucleus of the stria terminalis, the amygdala, and the dorsal hippocampus, because they are cerebral areas involved in cognition and emotional regulation, and they are also known to be affected by alcohol consumption [33, 34]. Moreover, the immunomarkers included in this study are highly expressed in these brain regions [35]. Moreover, we wanted to investigate these objectives at two ages: late adolescence and adulthood. Craving was also assessed, and the time points chosen (acute: 24h; early: seven days; protracted: 28 days) were selected, based on previous studies that reported the emergence of these negative affective behaviours [23, 36, 37]. The potential impact of these alterations could constitute an early index of neuroadaptation in alcohol dependence.

## 2. MATERIALS AND METHODS

### 2.1. Animals

Forty-two male C57BL/6J mice were acquired from Janvier Labs (Le Genest-Saint-Isle, France) on postnatal day (pd) 21. The mice were single-housed in cages containing nesting material, water and food provided *ad libitum*, in a 12h light-dark cycle (lights on at 8:00 a.m.) at standard humidity and temperature conditions. To differentiate between the age of adolescence and adulthood, objective criteria were used [38]. The procedures were in accordance with the European Directive 2010/63/UE, the National Institutes of Health guide for the care and use of laboratory animals (NIH Publications No. 8023, revised 1978) and Spanish regulations (Real Decreto 53/20130 and Ley 32/2007) for animal research. The experimental protocols were approved by the research ethics committee of the University of Oviedo and the Council of Rural Areas and Territorial Cohesion of the Principality of Asturias (code: PROAE 09/2021).

### 2.2. Experimental design

The mice were randomly assigned to control cage (CON, n = 12), water + behaviour (Water, n = 14) and ethanol + behaviour (EtOH, n = 16) conditions. The animals’ body weight and welfare were evaluated weekly, and all efforts were made to minimise animal suffering and reduce the number of animals used (supplementary material). As shown in Fig. 1, a two-bottle choice procedure was carried out as a model of voluntary alcohol consumption [39, 40]. Over five weeks, the EtOH group was given 24-h concurrent access to one bottle of alcohol in tap water and one bottle of water for four consecutive days, followed by three days of alcohol deprivation (on which only water was available) [41, 42, 43]. On the fourth day of each week, we placed lickometers in each bottle to register the numbers of licks made by the animals, in order to obtain a more accurate measure about the consumption (the procedures and results are displayed in the supplementary section). The EtOH concentration increased over the weeks (1º week: 3%; 2º week: 6%; 3º week: 10%; 4º and 5º weeks: 15%) and it was unsweetened. Water and alcohol consumption were recorded daily at 8:00 a.m. Fresh water and alcohol solutions were prepared daily, and the positions of bottles in the cage were switched daily to avoid any side bias. Throughout the experiment, two bottles containing water and alcohol in a cage without mice were used to evaluate the spillage/evaporation due to the experimental manipulations. Following the same schedule, two bottles containing water were offered to the CON and Water groups.

**Fig. 1.**
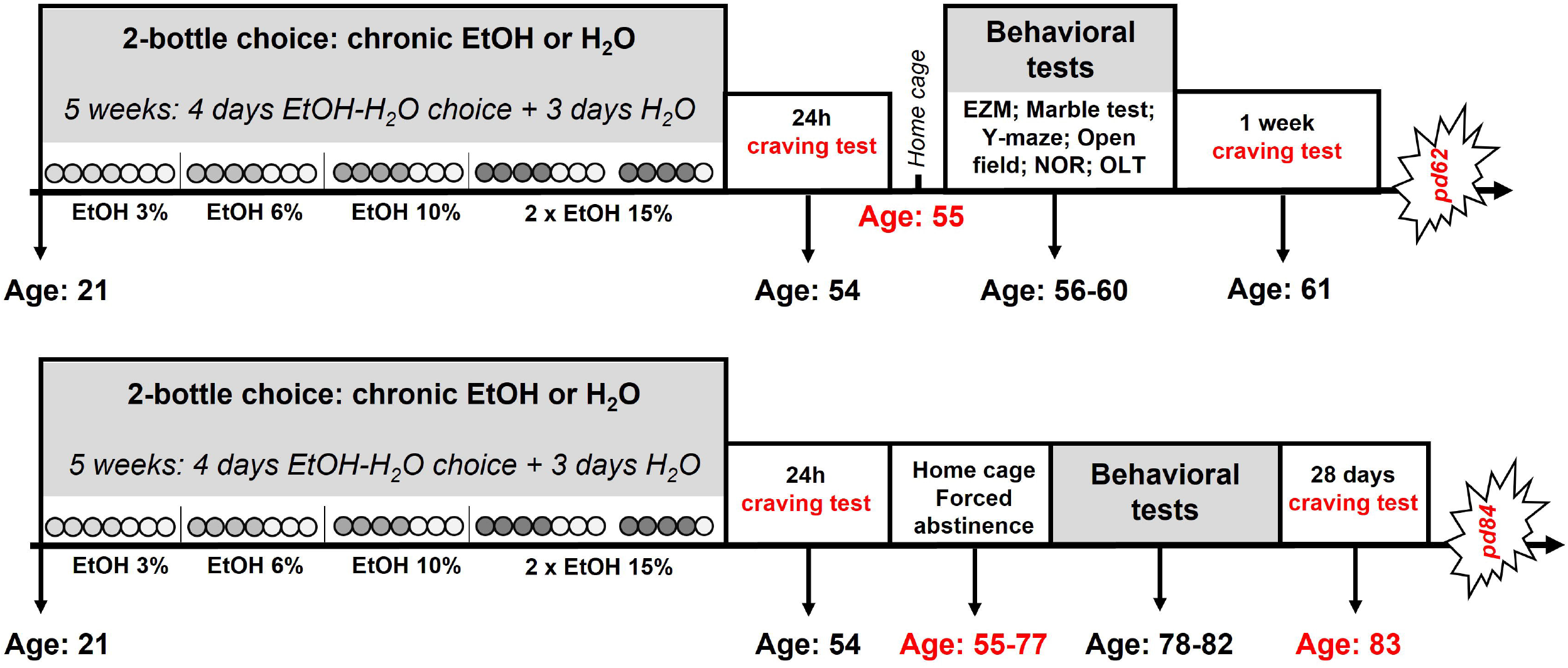
Experimental design. Mice were exposed to a two-bottle choice protocol for four days per week during 24h for five weeks (from pd21 to 52). Ethanol concentrations increased over the weeks, and in the fifth week remained at 15%. After 24h of abstinence, all animals were exposed again to two bottles of water and 15% alcohol to measure craving. Then, half of the animals were tested in different behavioural tests to study the impact of alcohol use in late adolescence on anxiety and cognition. After seven days, craving was assessed again, and mice were perfused the following day. The rest of the animals were maintained undisturbed under forced ethanol withdrawal in order to assess its impact in adulthood. During pd78-82, behavioural assessment was carried out and craving was analysed at pd83. These animals were perfused at pd84.

After the adolescent EtOH exposure (pd21-52), the EtOH group had a day off in the home cage and then, on pd54, underwent a 24h craving test in which two bottles containing water or 15% EtOH were offered to the animals over 24h. On pd55, the EtOH group had a rest day in the home cage before starting the behavioural assessment. Hence, we avoided the possible negative effect of the alcohol intake and the handling of the tests. Half of the animals in the EtOH (n=8) and Water (n=7) groups performed a battery of behavioural tests (pd56-60), while the other half were kept in their home box under forced abstinence. When the groups were randomly assigned, we verified that there were no differences in alcohol preference in the fifth week of the two-bottle choice paradigm (*t*_(14)_=0.32, *p*=0.75). After seven days (pd61) from the last experience with alcohol, the animals that performed the behavioural tests had another craving test, and 24h afterwards, the EtOH group was sacrificed, together with six animals from the CON group (pd62; Fig.1). The remaining mice were kept in their home boxes (pd55-77) and performed the same behavioural tests (pd78-82). On pd83, alcohol craving was assessed (after 28 days of forced abstinence) and the following day (pd84), animals were sacrificed (six from the CON group, eight from the EtOH group and seven from the Water group) (Fig.1).

In these craving tests, after 24h (short withdrawal), one week (intermediate withdrawal) and 28 days (protracted withdrawal) from the last alcohol experience, we presented two bottles with 15% EtOH and water during 24h and calculated the animal consumption (g/kg and preference).

### 2.3. Behavioural assessment

The behavioural tests (Fig.1) were performed based on previously published protocols from days 56-62 for mice in late adolescence and from days 78-84 for animals in adulthood [44, 45]. This behavioural battery of tests allowed the assessment of exploratory activity, anxiety-related behaviours and cognitive domains in the animals. Briefly, for an evaluation of compulsive and anxiety-like behaviours, the mice performed one session in the elevated-zero maze (EZM) and the marble burying test (MBt) (day 56 for adolescent mice and day 78 for adult animals). To assess spontaneous alternation as well as short-term memory, the mice underwent the *Y*-maze test (day 57 for adolescent mice and day 79 for adult animals). The following day (day 58 for adolescent mice and day 80 for adult animals), one session in the open field (OF) was carried out to analyse exploratory-related behaviours and, in the same environment, but 3h later, two identical copies of an object were placed near two adjacent corners, and mice were left to explore for 10 min (sample session). On day 59 for adolescent mice and day 81 for adult animals, the animals were allowed to explore for 10 min an identical copy of the familiar object and a ‘novel’ unknown object, located in the previous positions (novel object recognition test (NOR)). Finally, on Day 60 for adolescent mice and day 82 for adult animals, the mice explored for 10 min two identical copies of the familiar object: one of them placed in its habitual position, and the other displaced to an opposite corner (object location test (OLT)). Behavioural variables were analysed automatically, some of them manually, with Ethovision XT 12 software (Noldus, Wageningen, The Netherlands). Details about the apparatus’ dimensions and behavioural variables registered can be reviewed in the Supplementary Material section.

### 2.4. Immunohistochemistry and microscopy

On days 62 and 84, the mice were intracardially perfused for brain tissue processing with 0.1M phosphate-buffered saline, pH 7.4 (PBS), and 4% paraformaldehyde solution in PBS. Their brains were post-fixed for 24h in paraformaldehyde at 4ºC, after which they were dissected through the interhemispheric fissure. The right hemisphere (which was arbitrarily chosen to perform the immunohistochemical procedures) was cut into coronal sections (40 μm) using a vibratome (MICROM HM 650V, Thermo Scientific) and following a 1/6 serialisation (every sixth brain slice was analysed, leaving five sections unevaluated). The sections first received a heat-induced epitope retrieval (EnVision Flex high pH solution; Dako, Glostrup, Denmark), followed by an endogenous peroxidase blocking (80% PBS, 10% alcohol and 10% hydrogen peroxidase) for 30 min in the dark. To investigate the effect of EtOH consumption on brain mechanisms involved in HPA axis functioning and brain plasticity, we studied the expression of anti-glucocorticoid receptors (a polyclonal rabbit antibody at 1:200 (M-20), Santa Cruz Biotechnology), anti-CRF (a monoclonal rabbit at 1:100 (ab272391), Abcam) and anti-BDNF (a monoclonal mouse antibody at 1:100 (SAB4200744), Merck) primary antibodies. On the second day, the sections were incubated for 90 minutes in a biotin conjugated secondary antibody (a goat anti-rabbit, a goat anti-mouse or a swine anti-rabbit as appropriate; Dako; diluted 1:500) and for 1h in a peroxidase conjugated extravidin (Sigma, 1:1000 in PBS) solution in the dark. Diaminobenzidine (DAB) was employed as a chromogen with nickel at 0.04% in the case of CRF immunohistochemistry. Finally, the sections were dehydrated in alcohol, cleared in xylene and coverslipped with an EUKITT mounting medium. Negative controls in which the primary antibody was omitted resulted in absent staining.

The counting was conducted in one out of every six sections (1/6 serialisation), leaving five sections unevaluated, by an experimenter blind to the experimental condition. A total of four sections per animal were studied in regard to the cingulate, prelimbic and infralimbic cortices (Cg, PL, IL, 1.98 to 1.54 mm from bregma), the bed nucleus of the stria terminalis (BNST, 0.50 to 0.26 mm from bregma), the paraventricular nucleus of the hypothalamus (PVN, −0.70 to −1.06 mm from bregma), the central and the basolateral amygdala nuclei (CeA and BlA, −1.06 to −1.70 mm from bregma) and the dorsal hippocampus (dentate gyrus (DG) and cornu ammonis (CA3 or CA1), between −1.22 and −2.54 mm from bregma). For BDNF quantification, high-resolution photographs (10x) were captured with an Olympus BX51 microscope equipped with an Olympus DP70 digital camera (Olympus, Glostrup, Denmark). The area of each region of interest was drawn, measured, and the BDNF positive cells within the region were manually counted and expressed as their average number per unit area (mm^2^). In the case of GR and CRF immunohistochemistry, quantification was performed by densitometry. Background staining was controlled in each section by calculating the average optical density levels from another part of the section without labelling (10 measures), and subtracting this value from the average measure of each region. For this purpose, we used ImageJ software (US National Institutes of Health, Maryland, USA) [46, 47].

### 2.5. Statistical analysis

Data analysis was performed in Statistica 8 (StatSoft, Software Inc.). All data were reported as mean ± SEM, with *p* ≤ 0.05 as the criterion for statistical significance. To analyse the normal distribution of data, we performed a Shapiro-Wilk normality test. Alcohol consumption during 24h was calculated as grams of alcohol intake per kilogram of mouse body weight (g/kg), and preference as (ml EtOH/(ml EtOH + ml water))*100. Changes in alcohol consumption and preference over the weeks were assessed through repeated ANOVA measures. Comparisons among groups in behavioural tests were analysed by a two-way ANOVA (factors: age (late adolescence/adulthood) and EtOH (presence/absence)), while HSD Tukey’s post-hoc test was performed when differences were found (Fig.3). Expressions of GR, CRF and BDNF immunohistochemical markers were expressed as the percentage of change relative to the CON group ((animal score *100)/mean CON group). The percentage of change was calculated from the means of the basal group. After this correction, the data of EtOH and Water groups were analysed by an independent samples *t*-test (Fig.4).

## 3. RESULTS

### 3.1. Behavioural tests

#### Two-bottle choice drinking paradigm

EtOH consumption and preference during the two-bottle choice protocol increased over the sessions (*F*_(19,285)_=10.22, *p<*0.001; *F*_(19,285)_=2.16, *p=*0.004, respectively) (Fig.2a). Repeated ANOVA measures showed that there were significant differences in EtOH consumption during the last week of the two-bottle choice drinking paradigm (fifth week) and the abstinence sessions (*F*_(3,21)_=14.91, *p=*0.001), but not in the preference (*F*_(3,21)_=1.16, *p=*0.31). Specifically, Tukey’s post-hoc test revealed a higher EtOH consumption in the seven days (*p*=0.0002) and 28 days (*p*=0.002) craving tests respectively, to the fifth week. Besides this, the EtOH group consumed more EtOH in the craving test after seven days of alcohol withdrawal compared to 24h (*p*=0.005) (Fig.2b).

**Fig. 2.**
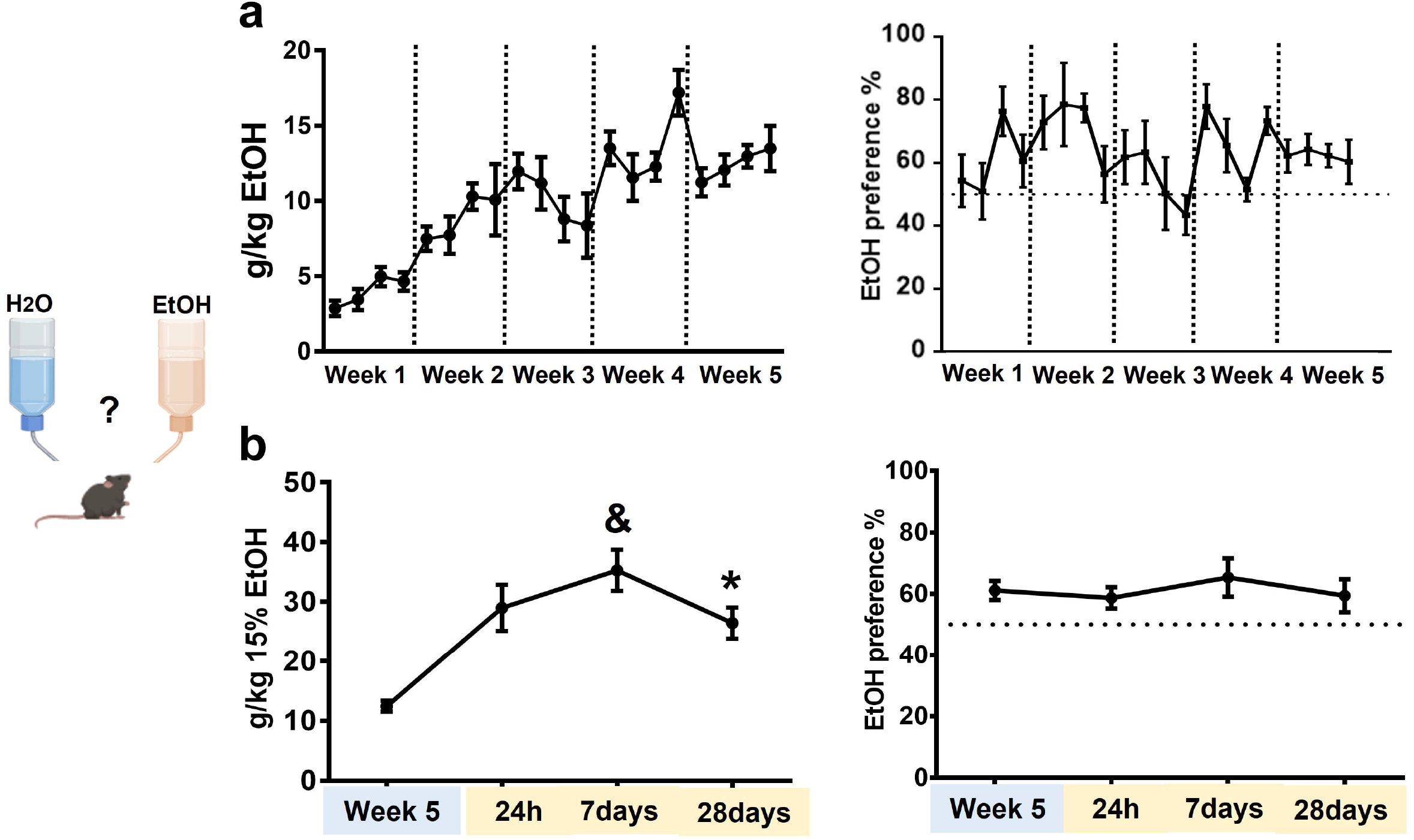
Changes in alcohol consumption and preference during a two-bottle choice protocol. EtOH consumption and preference during the two-bottle choice protocol increased over the sessions (*p*<0.001; *p*=0.004, respectively) (Fig.2a). There were significant differences in the EtOH consumption among the last week of the two-bottle choice drinking paradigm (fifth week) and the abstinence sessions (*p*=0.001), but not in the preference (*p*=0.31). Post-hoc analysis revealed a higher EtOH consumption in the seven days (&*p*=0.0002) and 28 days (\**p*=0.002) craving tests in respect to the fifth week. The EtOH group consumed more EtOH in the craving test after seven days of alcohol withdrawal compared to 24h (*p*=0.005) (Fig.2b).

#### EtOH consumption during early adolescence, increased anxiety, and compulsive-related behaviours in late adolescence and adulthood

EtOH consumption during early adolescence reduced the time spent in the open sections of the EZM (*F*_(1,26)_=4.28, *p*=0.05), as well as the indexes of time by entries and anxiety (*F*_(1,26)_=11.05, *p*=0.003; *F*_(1,26)_=4.28, *p*=0.05, respectively). The adult mice travelled longer distances than adolescent mice, regardless of the EtOH condition (*F*_(1,26)_=8.07, *p*=0.01). The marble burying test allowed us to assess repetitive and compulsive behaviour. The two-way ANOVA showed a significant difference in the EtOH factor, revealing that this condition increased the number of marbles buried (*F*_(1,26)_=4.28, *p*=0.05). Finally, in the OF test, the interaction EtOH x age was significant in the variables distance travelled (*F*_(1,26)_=4.83, *p*=0.04) and thigmotaxis (*F*_(1,26)_=23.47, *p*=0.0001), showing that adult mice with EtOH experience travelled longer distances on the periphery of the maze. On the other hand, EtOH animals, regardless of their age, spent more time performing grooming behaviour (*F*_(1,26)_=8.43, *p*=0.007). Finally, adult mice performed more rearing behaviour (*F*_(1,26)_=14.54, *p*=0.007) over a greater amount of time (*F*_(1,26)_=16.69, *p*=0.003), and showed higher latency to enter into the centre of the field (*F*_(1,26)_=6.68, *p*=0.02) (Fig. 3).

**Fig. 3.**
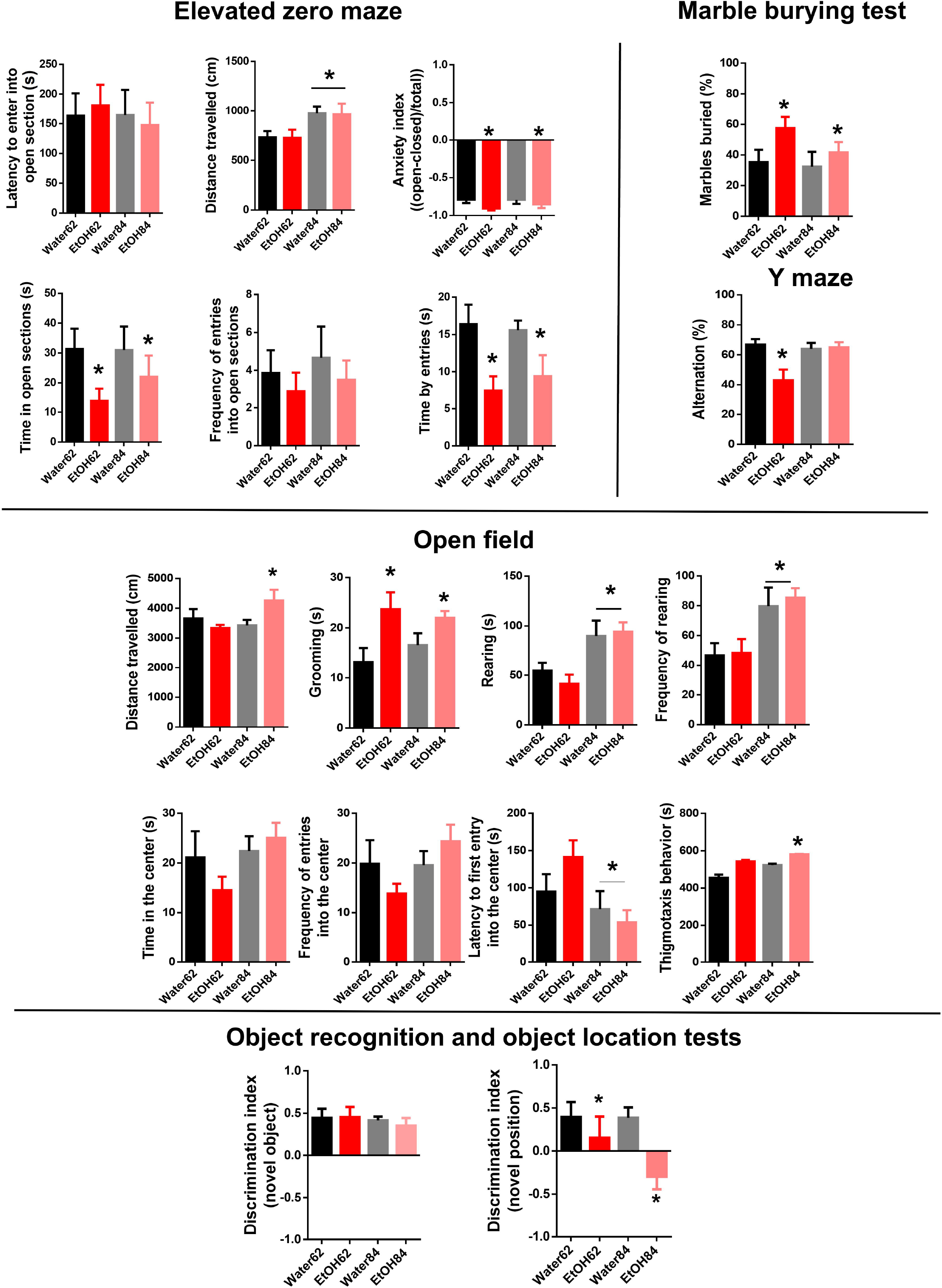
Behavioural performance of the control and EtOH groups during late adolescence and adulthood. On the one hand, anxiety, compulsive and exploratory-related behaviours were assessed through an elevated-zero maze test, a marble burying test and an open field. On the other hand, working memory, as well as of recognition of novel position and object, were tested by Y-maze, object location and novel object recognition tests. All data were mean ± S.E.M., and statistically significant differences were considered when *p*≤0.05 (*).

#### EtOH consumption during early adolescence altered working memory in late adolescent mice

*Y*–maze test performance was altered in late adolescent mice of the EtOH group (*F*_(1,26)_=6.40, *p*=0.02), but this effect did not ensue in the adult animals.

#### EtOH consumption during early adolescence worsened performance in the OLT, whereas NOR was unaltered

Differences between the groups were not observed during the time of sniffing the two identical objects in the sample trial (*p*>0.05). In contrast, alcohol during early adolescence negatively affected the recognition of a new object position (OLT: *F*_(1,26)_=6.68, *p*=0.02), but not recognition of a novel object (*p*>0.05) in both ages (Fig. 3).

### 3.2. Immunohistochemical markers

#### EtOH consumption during early adolescence diminished BDNF+ cells in the Cg prefrontal cortex, CA1, and CeA regardless of the animals’ age. Reduction in CA3 was specific for mice on pd62

Alcohol consumption reduced BDNF expression in the Cg cortex, CA1, and the CeA of EtOH drinking mice, regardless of their age (*F*_(1,26)_=6.61, *p*=0.02; *F*_(1,26)_=4.85, *p*=0.04; *F*_(1,26)_=4.02, *p*=0.05, respectively). In the hippocampal subregion CA3, the BDNF was reduced only in late adolescent mice (*F*_(1,26)_=4.72, *p*=0.04). Besides this, BDNF increased with age in the BlA (*F*_(1,26)_=16.11, *p*=0.0006), CA1 (*F*_(1,26)_=16.29, *p*=0.0005), and the PL cortex (*F*_(1,26)_=6.29, *p*=0.02), whereas in the IL cortex (*F*_(1,26)_=4.70, *p*=0.04) it decreased (Fig. 4 a, b, c).

**Fig. 4.**
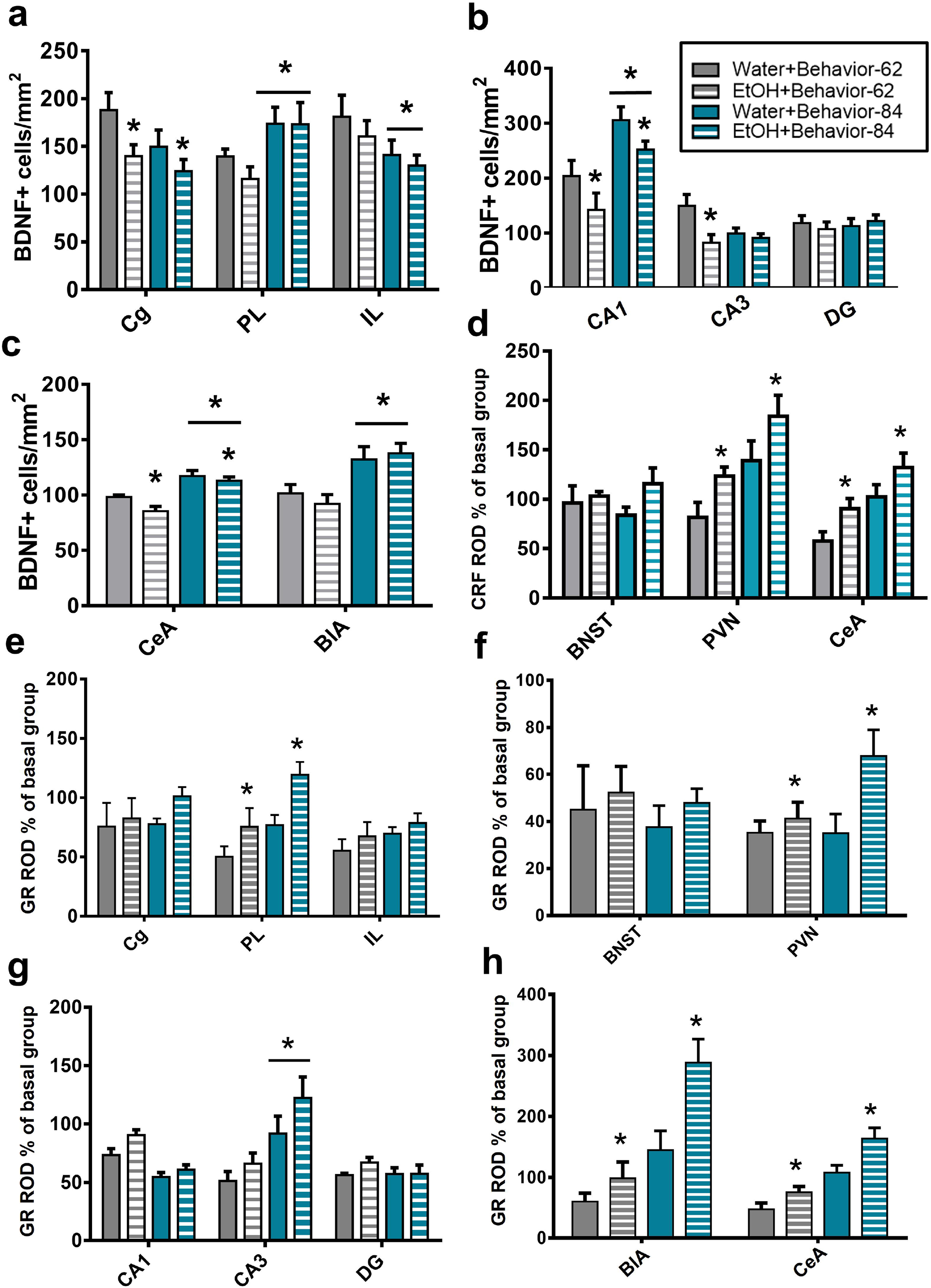
**(a, b, c)** EtOH consumption during early adolescence diminished BDNF+ cells in the Cg prefrontal cortex (\**p*=0.02), CA1 (\**p*=0.04) and CeA (*p*=0.05), regardless of the animals’ age. Reduction in CA3 was specific for mice on pd62 (\**p*=0.04). BDNF increased with age in the BlA (<u>*</u>*p*=0.0006), CA1 (<u>*</u>*p*=0.0005), and the PL cortex (<u>*</u>*p*=0.02), whereas in the IL cortex (<u>*</u>*p*=0.04) it decreased. **(d)** EtOH consumption during early adolescence enhanced CRF density in the PVN of the hypothalamus (\**p*=0.02) and the CeA (\**p*=0.02) **(e, f, g, h)**. EtOH consumption during early adolescence increased GR density in the PL prefrontal cortex (\**p*=0.0006), the PVN of the hypothalamus (\**p*=0.03), and both the CeA (\**p*=0.005) and the BlA nuclei (\**p*=0.007). Adult mice had higher GR density in the CA3 compared to late adolescent animals, regardless of their EtOH condition (*<u>*</u>p*=0.001). BDNF, CRF and GR expression are expressed as a percentage of change relative to the CON group ((animal score *100)/mean CON group). The results are represented as the mean number of positive cells per mm^2^ + S.E.M. Abbreviations: Cg: cingulate cortex; PL: prelimbic cortex; IL: infralimbic cortex; PVN: paraventricular nucleus; CA: cornu ammonis; DG: dentate gyrus; CeA: central amygdala; BlA: basolateral amygdala.

#### EtOH consumption during early adolescence enhanced CRF density in the PVN of the hypothalamus and the CeA, but not in the BNST

Interaction *age x EtOH* was significant in the PVN (*F*_(1,26)_=5.69, *p*=0.02), revealing differences between age groups only when the EtOH was present. CRF density was also increased in the CeA of the EtOH group (*F*_(1,26)_=6.75, *p*=0.02) in respect to the Water group, regardless of age. CRF density in the BNST did not show significant differences (*p*>0.05) (Fig.4 d).

#### EtOH consumption during early adolescence increased GR density in the PL prefrontal cortex, the PVN of the hypothalamus, and both the CeA and the BlA nuclei

GR density was higher in the PL cortex, the PVN, and both the CeA and the BlA nuclei of the EtOH group compared with the Water group, regardless of age (*F*_(1,26)_=8.94, *p*=0.0006; *F*_(1,26)_=5.32, *p*=0.03; *F*_(1,26)_=9.42, *p*=0.005; *F*_(1,26)_=8.45, *p*=0.007, respectively). Moreover, adult mice had higher GR density in the CA3 compared to late adolescent animals, regardless of the EtOH condition (*F*_(1,26)_=13.70, *p*=0.001) (Fig. 4 e, f, g, h).

Representative photographs of each marker for each brain region analysed are displayed in Figure 5. along with their Bregma levels.

**Fig. 5.**
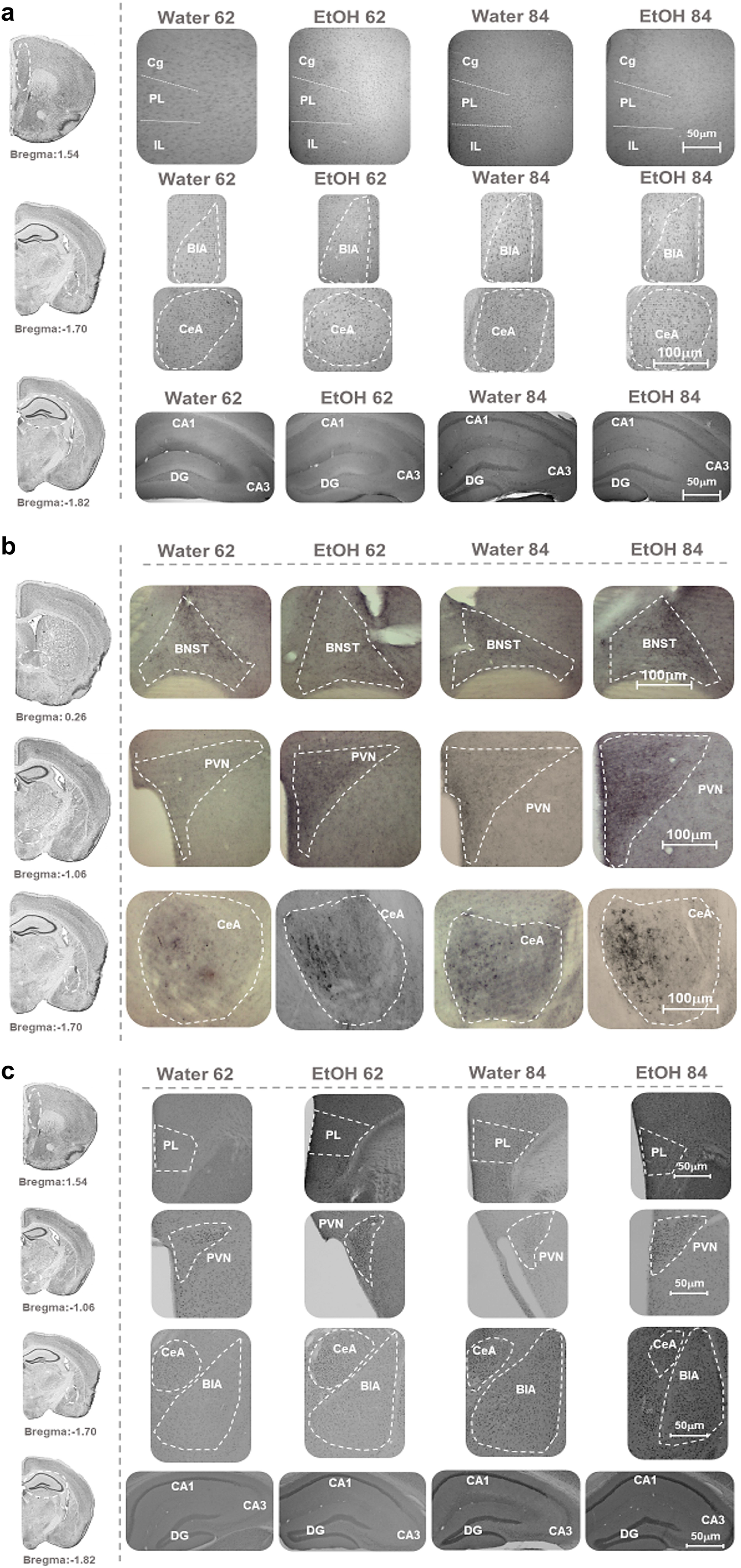
**(a)** Representative photographs of the areas quantified for BDNF (the brain hemispheres were separated, and only the right hemisphere is shown). Images were taken at 10X magnification. **(b)** Representative photographs of the areas quantified for CRF (only the right hemisphere is shown). Bregma references are indicated in the images, which were taken at 10X magnification. **(c)** Representative photographs of the areas quantified for GR (the right hemisphere is shown). Bregma references are shown in the images, which were acquired at 4X magnification [90].

## 4. DISCUSSION

The present study aimed to assess the emotional and cognitive alterations caused by alcohol consumption in adolescence at two time-points: late adolescence and adulthood. Due to the anxious and compulsive behaviours found in this research, we further analysed the expression of markers related to HPA axis activity at both ages. Finally, we were also interested in assessing craving after forced alcohol withdrawal at three points: 24h, seven and 28 days. Given the high prevalence of alcohol use during adolescence, it is important to understand the long-term consequences of adolescent ethanol exposure.

### Adolescent alcohol is associated with long-lasting emotional and cognitive alterations in mice

We used the two-bottle choice paradigm to model the pattern of voluntary consumption observed in humans because this procedure is technically simple, and it leads to levels of ethanol intake that are considered clinically relevant [48]. In our study, ethanol intake after withdrawal was assessed at acute, early and protracted abstinence, showing an increase after seven and 28 days compared to the last two-bottle choice session. This result could be explained by a different hypothesis related to negative reinforcing effects of the alcohol, as well as an effect facilitated by intermittent vs. continuous chronic ethanol exposure. Since we observed enhanced withdrawal-related anxiety-like behaviour in the EZM test, compulsivity in the marble burying test, more grooming behaviour in the OF test, as well as memory problems in the *Y*-maze test and in the OLT, it seems that a negative effect triggered by abstinence could have induced a higher ethanol consumption. These results are consistent with previous studies that also observed anxiety-like behavioural alterations, which may be involved in perpetuating this pattern of alcohol abuse [49, 50]. Unfortunately, some of the emotional and cognitive alterations found were maintained into adulthood, even during a long period of abstinence, suggesting the need for interventions at an early age to avoid later alcohol dependence. In this sense, previous studies of our group have found a positive effect from aerobic exercise and cognitive training, which are valuable strategies to counteract some behavioural alterations associated with ethanol intake during adolescence and early adulthood [44, 51].

Interestingly, performance in the NOR test was preserved in both ages, whereas an altered discrimination index was found in the OLT. While both tests assessed the inherent preference of mice for novelty to reveal memory for previously explored objects, the OLT primarily evaluates spatial learning, which relies heavily on hippocampal activity, while the NOR test evaluates non-spatial learning, which relies on multiple brain regions [52]. Although some studies suggest that ethanol does not affect spatial learning tasks [53, 54], other researchers have shown that ethanol use selectively impairs spatial hippocampal dependent memory in animals [55, 56]. According to our results, a classic study [57] has also found that acquisition and subsequent retention of spatial memory is more impaired in adolescent rats compared with adults, and the acquisition of non-spatial learning was not affected by ethanol in rats of either age. Regarding this, ethanol intake has been shown to reduce hippocampal neurogenesis and inhibit several hippocampal synaptic plasticity mechanisms underlying memory [58, 59, 60].

Meanwhile, performance in the *Y*-maze test employed to assess spatial working memory was only altered in mice during late adolescence, but not in adulthood. Preserved prefrontal cortical functions were required to carry out correctly spontaneous alternation [61], suggesting that protracted ethanol abstinence could have reserved this deficit. However, other studies have found that working memory deficits in more challenging tasks, such as the radial-arm water maze, did not recover after months of ethanol abstinence [62, 63].

### Adolescent alcohol is associated with changes in stress-related markers (BDNF, GR and CRF) in different brain regions

Similar to stress, alcohol intake activates the HPA axis to release cortisol in humans and corticosterone (CORT) in rodents from the adrenal gland [64], contributing to the reinforcing effects of the drug [65, 66]. The consequences of this dysregulation for alcohol abuse and relapse remain unclear. However, and owing to our behavioural results mentioned above, we suppose that under abstinence, alcohol is used to alleviate or prevent negative emotional symptoms, such as anxiety, compulsivity or anhedonia, which emerge in the absence of the drug [67].

In our study, we observed a higher GRs density in brain regions such as the PL cortex, the PVN, as well as the central and basolateral amygdala nuclei in both late adolescent and adult mice exposed to alcohol during early ages. According to this finding, it seems that alcohol consumption, and above all, abstinence, induce an upregulation of GRs levels in stress/reward-related brain regions [68]. Thus, although peak HPA axis activation is blunted during alcohol withdrawal, the GRs dysregulation seems to remain [69]. Hence, a sensitized GR system would be expected to sustain the escalation of alcohol intake even in the absence of peak levels of released CORT. On the other hand, and regardless of age, our EtOH mice also expressed higher levels of CRF in brain regions such as the PVN and the CeA, both characterised by high expression of this marker. The function of CRF in the PVN is essential for activation of the HPA axis, and the CeA is involved in stress and addiction-like behaviours regulating the stress response and peripheral processes involved in somatic signs of withdrawal [70, 71, 72]. In particular, the CeA is a key brain region with - and interface role between - stress and addiction-related processes [73]. Furthermore, injections of CRF and CRF receptor agonists exacerbated alcohol withdrawal-induced stress [74], increased anxiety-like behaviour in the elevated-plus maze [75], enhanced withdrawal-induced social anxiety and increased the footshock stress-induced reinstatement of alcohol seeking [76]. Thus, this evidence could suggest that alcohol intake and relapse could be reduced by CRF manipulations [73].

Stress is also known to influence BDNF expression, and this neurotrophic factor has been implicated in several stress-related neuropsychiatric disorders, including alcohol addiction [77]. Preclinical studies have revealed that chronic exposure to alcohol leads to depressive and anxious-like behaviours and a decrease in the central expression of BDNF, these remaining the underlying unknown neurobiological mechanisms [78, 79]. In our study, we observed a reduction of BDNF expression in the Cg cortex, the CA1, and the CeA of the EtOH groups (pd62 and 84). In general, both acute and chronic stress experiences have been shown to reduce BDNF expression in these same brain areas (prefrontal cortex [80, 81], central amygdala [82], and hippocampus [83]). Further, accumulating evidence indicates cross talk between GRs and BDNF systems [30, 84], and optimal functioning of the BDNF-TrkB-ERK signalling may be a susceptibility factor for developing memory impairments and altered hippocampal functions, which is consistent with our behavioural results in the OLT [85].

We would like to mention that the *age* factor was significant, showing an age-related increase of BDNF and GRs expression in some brain regions of pd84 mice compared with pd62. Throughout prenatal and early postnatal development, BDNF is expressed at very low levels, but it undergoes a rapid increase in expression at around four weeks of age, followed by a decline over the mice’s lifespan [86]. On the other hand, adolescent rats and mice exposed to physical and/or psychological stressors often exhibit ACTH and corticosterone responses that last twice as long as those observed in adults [87, 88], which could affect GRs expression involved in the negative feedback regulation of the HPA axis. These age-related changes, probably in response to variations in circulating gonadal hormones, extend the interest in these markers for understanding differences in susceptibility to substance dependence. Finally, and although it is not part of the aim of this study, we would like to mention the need to consider sex-related differences in the addiction research field. Thus, in several experiments there have been found sex and gender differences in addiction and relapse, both in humans and in animal models [89]. We cannot directly extrapolate our result to females, and for this reason, more studies must be conducted in order to elucidate this issue.

## 5. CONCLUSIONS

In conclusion, our study revealed that ethanol consumption during early years is related to several emotional and cognitive alterations, of which some of them remained in adulthood. We also observed that forced ethanol withdrawal induces an abstinence effect at different time points, which were higher at seven days. Moreover, these behavioural results could be related, in part, to the neurobiological changes observed in the expression of BDNF, as well as in immunohistochemical markers involved in HPA axis functioning (GR and CRF) after alcohol use. Further research could be aimed at determining the temporal dynamics of the observed alterations in BDNF, GR and CRF expression at different time points of ethanol withdrawal, as well as its relationship with emotional and cognitive changes.

## Acknowledgements

We thank Claudia Jove and Pablo Piñera for their help during behavioural procedures.

## Authorship contribution statement

Patricia Sampedro-Piquero: conceptualisation, methodology, validation, formal analysis, investigation, data curation, visualisation, writing - original draft, writing - review & editing. Román D. Moreno-Fernández: methodology, investigation, data curation, visualisation, writing - review & editing. Azucena Begega: resources, funding acquisition; Matias López: conceptualisation, funding acquisition. Luis. J. Santín: data curation, resources, writing - review & editing. All the authors have approved the final version of the manuscript.

## Data availability statement

All relevant data are presented in the manuscript; raw data is available upon request from the corresponding author.

