## Supplementary material for "Long-term consequences of alcohol use in early adolescent mice: focus on neuroadaptations in GR, CRF and BDNF"

Animal's body weight was evaluated weekly as shown in the following figure (Figure 1). Body weight increased over the weeks ($F_{(8,192)}$=729,60, $p$<0.001).

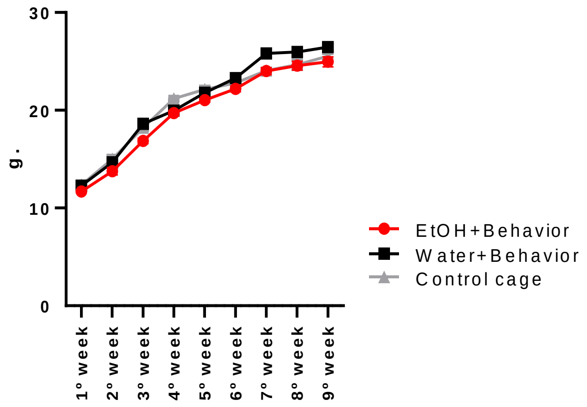

Figure 1. Body-weight gain across the weeks in the different experimental conditions.

### Supplementary methods for 2-bottle choice protocol

The last day of each week, as well as in the abstinence tests (24h, 7 and 28 days), a contact sensitive lickometers registered the licks made by mice to the nearest 0.01 s, and MED-PC software (Med Associates, Inc) controlled the equipment and recorded the data.

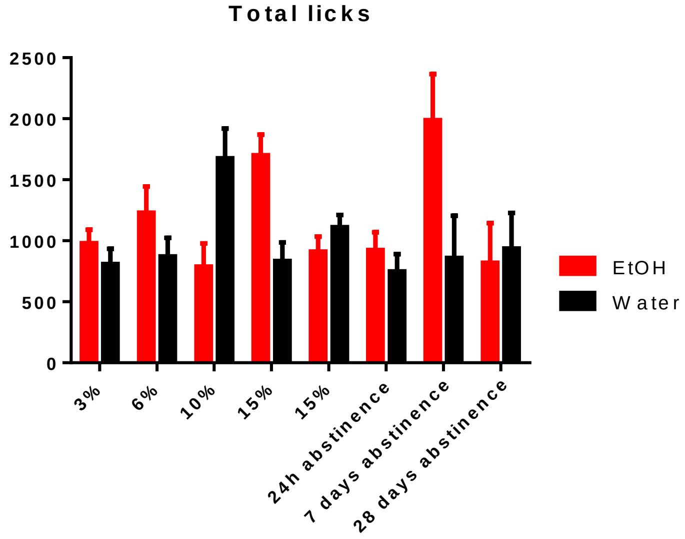

**Supplementary methods for the behavioral assessment**

Behavioral procedures were conducted between 9:00 a.m. and 3:00 p.m. in noise-isolated rooms illuminated 50 lux. Mice were daily habituated to the testing room for at least 20 minutes before starting behavioral assessment. Sessions were videotaped with an overhead digital camera and analyzed with the software Ethovision XT (Noldus, Wageningen, The Netherlands). After each session, a 10% alcohol solution was used to clean the arena of the dry apparatuses and eliminate odor cues.

Elevated-zero maze (EZM)

EZM apparatus was raised 50 cm above the floor and consisted of two open and two closed sections (wide 5 cm). The inside diameter was 35 cm. Test time was 5 minutes. Latency to enter in the open section (s), distance travelled (cm), time in open sections (s) and frequency of entries into open sections were registered. We also calculated two indexes: anxiety index (time in open sections-time in closed sections/total) and time by entries index (time spent in the open section/square-root of number of entries for each animal). Therefore, high TbE rating is indicative of low levels of anxiety, while low TbE rating is indicative of high levels of anxiety [1, 2].

Marble-burying test

This test was carried out in a standard laboratory cage for each mouse (20x35x55 cm). Testing was conducted under dim lighting (20 lx) and the mouse was placed in a new, clean cage with a bedding thickness of 5 cm and allowed to freely dig for 10 min (without marbles). The mouse was then removed from the cage. The bedding was flattened and 15 marbles were arranged in a 5 × 3 array on top of the bedding (Figure 2). The mouse was reintroduced to the cage and allowed to bury the marbles for 30 min. The number of marbles that were buried (covered two-thirds or more by bedding) was counted at the end of the test by an observer who was blind to treatment group [3].

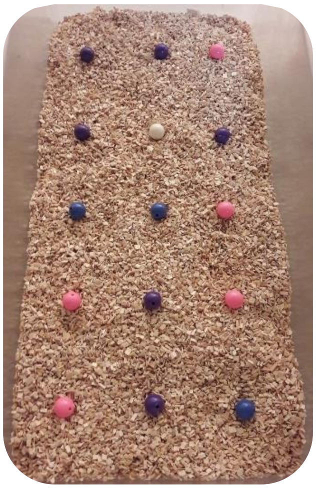

Figure 2. Disposition of the marbles in the test.

Y maze

Rodents tend to explore the least recently visited arm alternating the visits between the three arms. Hence, working memory is necessary to maintain an ongoing record of most recently visited arms, and continuously update such record. The maze consisted of three equally sized arms (30 x 14 cm each), with opaque walls (33 cm high). Test time was 8 minutes [4].

Open field – NOR – OLT

Mice were allowed to explore a squared open field (42 x 42 cm; 40 cm high) with a central area considered as a 20 x 20 cm square. Test time was 10 minutes. Distance travelled, time of grooming and rearing behaviors (s), frequency of rearing, time in the center (s), frequency of entries into the center, latency to first entry into the center (s), time exploring the periphery of the maze (thigmotaxis behavior) (s) [4,5].

The NOR and OLT tests were performance in the same field. Both tests lasted 5 minutes. In the NOR test (performed 3 hours after the habituation session), two identical copies of an object (i.e. 'familiar' object) were placed near two adjacent corners and mice were left to explore for 10 min (sample session). Next day, mice were allowed to explore for 10 min an identical copy of the familiar object and a 'novel' unknown object, located in the previous positions. 24h later, mice explored for 10 min two identical copies of the familiar object, one of them placed in its habitual position, and the other displaced to an opposite corner (OLT) (Figure 3). The total time of object exploration (defined as the mouse actively touching an object with its nose (sniffing) was scored by a trained observer. From this behavioral measurement, we calculated two ratio: a NOR ratio [(time exploring the novel object−time exploring the familiar object)/total time exploring both objects] and an OLT ratio [(time exploring the displaced object−time exploring the static object)/total time exploring both objects]. A positive ratio score that is significantly different from zero would indicate a successful object or place memory [6].

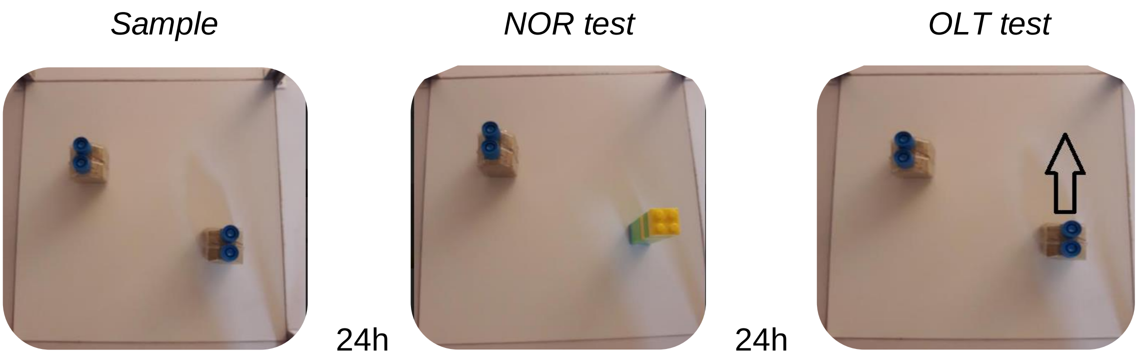

Figure 3. Schedule of the sample, object and place memory test.

### Supplementary methods for Immunohistochemistry

| ***Brain region*** | ***Abbreviation*** | ***Bregma levels*** |
|---|---|---|
| **Prefrontal cortex** | | |
| -Cingulate cortex | Cg | 1.98 to 1.54 mm |
| -Prelimbic cortex | PL | 1.98 to 1.54 mm |
| -Infralimbic cortex | IL | 1.98 to 1.54 mm |
| **Bed nucleus of the stria terminalis** | BNST | 0.50 to 0.26 mm |
| **Hypothalamus** | | |
| -Paraventricular nucleus | PVN | −0.70 to −1.06 mm |
| **Amygdala** | | |
| -Central nucleus | CeA | −1.06 to −1.70 mm |
| -Basolateral nucleus | BlA | −1.06 to −1.70 mm |
| **Dorsal Hippocampus** | | |
| -Cornu Ammonis 1 | CA1 | -1.22 and -2.54 mm |
| -Cornu Ammonis 3 | CA3 | -1.22 and -2.54 mm |
| -Dentate Gyrus | DG | -1.22 and -2.54 mm |

### Supplementary statistical analysis

Those behavioral variables registered without significant differences between groups will be shown below. Two-way ANOVAs, in which Age (pd62/pd84) and EtOH (ethanol vs. no ethanol) were the two factors with two levels each, were carried out:

*-Behavioral results*

| ***Test*** | ***Variable*** | ***F, p*** |
|---|---|---|
| **EZM** | Latency to enter in open sections (s) | EtOH: $F_{(1,26)}$=0.001, $p$=0.97<br>Age: $F_{(1,26)}$=0.22, $p$=0.64<br>EtOH x Age: $F_{(1,26)}$=0.20, $p$=0.66 |
| | Frequency of entries into open sections | EtOH: $F_{(1,26)}$=0.58, $p$=0.45<br>Age: $F_{(1,26)}$=0.20, $p$=0.65<br>EtOH x Age: $F_{(1,26)}$=0.01, $p$=0.93 |
| ***Y* maze** | Distance travelled (cm) | EtOH: $F_{(1,26)}$=0.38, $p$=0.54<br>Age: $F_{(1,26)}$=0.37, $p$=0.55<br>EtOH x Age: $F_{(1,26)}$=0.11, $p$=0.74 |
| **Open field** | Time in the center (c) | EtOH: $F_{(1,26)}$=0.31, $p$=0.58<br>Age: $F_{(1,26)}$=2.75, $p$=0.11<br>EtOH x Age: $F_{(1,26)}$=1.68, $p$=0.20 |
| | Frequency of entries into the center | EtOH: $F_{(1,26)}$=0.03, $p$=0.86<br>Age: $F_{(1,26)}$=2.40, $p$=0.13<br>EtOH x Age: $F_{(1,26)}$=1.68, $p$=0.21 |
| **NOR/OLT (Sample trial)** | Distance travelled (cm) | EtOH: $F_{(1,26)}$=0.12, $p$=0.66<br>Age: $F_{(1,26)}$=2.70, $p$=0.11<br>EtOH x Age: $F_{(1,26)}$=2.01, $p$=0.18 |
| | Sniffing both identical objects (s) | EtOH: $F_{(2,25)}$=1.13, $p$=0.86<br>Age: $F_{(2,25)}$=1.42, $p$=0.77<br>EtOH x Age: $F_{(2,25)}$=1.18, $p$=0.85 |

#### -Immunohistochemistry results

| ***Marker*** | ***Brain region*** | ***F, p*** |
|---|---|---|
| **BDNF** | **DG** | EtOH: $F_{(1,26)}$=0.01, $p$=0.94<br>Age: $F_{(1,26)}$=0.11, $p$=0.74<br>EtOH x Age: $F_{(1,26)}$=0.58, $p$=0.45 |
| **GR** | **IL** | EtOH: $F_{(1,26)}$=1.34, $p$=0.26<br>Age: $F_{(1,26)}$=1.95, $p$=0.18<br>EtOH x Age: $F_{(1,26)}$=0.03, $p$=0.87 |
| | **BNST** | EtOH: $F_{(1,26)}$=0.52, $p$=0.48<br>Age: $F_{(1,26)}$=0.24, $p$=0.63<br>EtOH x Age: $F_{(1,26)}$=0.01, $p$=0.90 |
| | **DG** | EtOH: $F_{(1,26)}$=1.04, $p$=0.32<br>Age: $F_{(1,26)}$=0.70, $p$=0.41<br>EtOH x Age: $F_{(1,26)}$=1.13, $p$=0.39 |
| **CRH** | **BNST** | EtOH: $F_{(1,26)}$=2.37, $p$=0.14<br>Age: $F_{(1,26)}$=0.1, $p$=0.99<br>EtOH x Age: $F_{(1,26)}$=0.97, $p$=0.33 |
